# Virus-like particle (VLP)-based vaccine targeting tau phosphorylated at Ser396/Ser404 (PHF1) site outperforms phosphorylated S199/S202 (AT8) site in reducing tau pathology and restoring cognitive deficits in the rTg4510 mouse model of tauopathy

**DOI:** 10.1101/2024.04.05.588338

**Authors:** Jonathan Hulse, Nicole Maphis, Julianne Peabody, Bryce Chackerian, Kiran Bhaskar

## Abstract

Tauopathies, including Alzheimer’s disease (AD) and Frontotemporal Dementia (FTD), are histopathologically defined by the aggregation of hyperphosphorylated pathological tau (pTau) as neurofibrillary tangles in the brain. Site-specific phosphorylation of tau occurs early in the disease process and correlates with progressive cognitive decline, thus serving as targetable pathological epitopes for immunotherapeutic development. Previously, we developed a vaccine (Qβ-pT181) displaying phosphorylated Thr181 tau peptides on the surface of a Qβ bacteriophage virus-like particle (VLP) that induced robust antibody responses, cleared pathological tau, and rescued memory deficits in a transgenic mouse model of tauopathy. Here we report the characterization and comparison of two additional Qβ VLP-based vaccines targeting the dual phosphorylation sites Ser199/Ser202 (Qβ-AT8) and Ser396/Ser404 (Qβ-PHF1). Both Qβ-AT8 and Qβ-PHF1 vaccines elicited high-titer antibody responses against their pTau epitopes. However, only Qβ-PHF1 rescued cognitive deficits, reduced soluble and insoluble pathological tau, and reactive microgliosis in a 4-month rTg4510 model of FTD. Both sera from Qβ-AT8 and Qβ-PHF1 vaccinated mice were specifically reactive to tau pathology in human AD post-mortem brain sections. These studies further support the use of VLP-based immunotherapies to target pTau in AD and related tauopathies and provide potential insight into the clinical efficacy of various pTau epitopes in the development of immunotherapeutics.

## Introduction

Aggregation of the microtubule associated protein tau (MAPT) into paired helical filaments (PHFs) and neurofibrillary tangles (NFTs) is one of the key pathological hallmarks of Alzheimer’s disease (AD) and related primary tauopathies.^1–3^ Post-translational modifications (PTMs) of the tau protein, mainly hyperphosphorylation, drives the pathological misfolding and aggregation process.^4,5^ The accumulation of tau aggregates in the brain is strongly associated with cognitive decline and cortical atrophy in AD indicating that tau pathology is one of the main drivers of the neurodegenerative process.^6–10^ Pathological tau (pTau) is also known to seed and spread trans-neuronally and seed further tau aggregation in a prion-like fashion.^11–19^ Based on this, it is important to develop therapeutics that specifically target pTau, promote its efficient clearance and thus prevent it from seeding/spreading in the brain.

Recent success in anti-amyloidβ antibody therapeutics such as Lecanemab^20^ and Donanemab^21^ have highlighted the efficacy of targeted antibody therapies at clearing protein aggregates from the brain. However, monoclonal antibody-based passive immunotherapies have major limitations including expensive manufacturing and consumer costs, the need for frequent and repetitive dosing to maintain therapeutic antibody levels, and the accessibility of infusion centers for drug administration.^22–26^ Active immunization using vaccines has the potential to avoid all of these caveats by harnessing the body’s immune system to generate a targeted antibody response against a desired immunogen. Virus-like particles (VLPs) are a highly effective approach for generating strong and durable antibody responses that can even overcome self-tolerance mechanisms. VLP-based vaccines have already been Food and Drug Administration (FDA) approved for the prevention of numerous infectious diseases,^27^ and are being investigated as vaccine platforms for numerous other disease conditions including AD.^28,29^ Qβ VLPs are recombinantly expressed Qβ bacteriophage coat proteins that spontaneously assemble into protein capsids that lack the genetic material to be infectious or replicate but strongly stimulate B cells.^30^ VLPs can thus serve as a platform to display antigens of interest on their surface in a highly immunogenic fashion owing to their dense, repetitive, and highly-ordered array on the VLP surface.^31,32^

Previously, we evaluated the efficacy of Qβ VLPs conjugated to phosphorylated tau at threonine 181 (Qβ-pT181) in the rTg4510 mouse model of human tauopathy. Qβ-pT181-immunized mice exhibited robust antibody responses, reduction in tau pathology, neuroinflammation, neurodegeneration and cognitive decline.^29^ However, tau protein can be differentially phosphorylated in AD and at different disease stages,^33^ therefore, it is possible that vaccines targeting a single epitope may be inadequate to modify the disease process for all patients at all disease stages. With this in mind, we sought to utilize the Qβ VLP platform to target other key pathologically modified epitopes of tau to address our central question: does the pTau epitope used as the immunogen in a Qβ VLP vaccine strategy impact the vaccine efficacy *in vivo*? For this study, we developed and tested two Qβ VLPs targeting the dual phospho-serine 199/phospho-serine 202 (pS199/pS202) epitope and the dual phospho-serine 396/phospho-serine 404 (pS396/pS404) epitope which we will refer to as the AT8 and PHF1 sites respectively based on the recognition sites for these two well-known phospho-tau antibodies.

The phosphorylated tau epitopes recognized by the antibodies AT8 and PHF1 have been implicated as earlier neuropathological accumulations in AD that correspond with intracellular and extracellular NFT formation.^33–39^ The phosphorylation sites recognized by AT8 (S199, S202, T205, and S208)^40–42^ and the phosphorylation sites recognized by PHF1 (S396 and S404)^43^ increase with disease progression in the brain.^36,37,44–46^ The S199 site in particular has shown dramatic increases in phosphorylation at later stages of AD compared to other tau residues.^36,44,45^ These sites may also reflect important steps in tau misfolding, assembly, or seed-competency contributing to the pathological process.^47–51^ The AT8 antibody has also been used as the primary post-mortem diagnostic tool for Braak staging, the neuropathological grading assessment for tau pathology in AD.^52^

We sought to evaluate the efficacy of Qβ-AT8 and Qβ-PHF1 vaccines in the rTg4510 mouse model overexpressing human mutant P301L tau.^53,54^ We observed that both vaccines successfully elicited robust IgG antibody responses against their respective pTau epitope but exhibited differential responses in protection against cognitive decline and tau pathology. The Qβ-PHF1 vaccine exhibited some protection against cognitive deficits, reductions in tau pathology with a preferential reduction of insoluble tau aggregates, and blunted microgliosis. In contrast, the Qβ-AT8 vaccine failed to demonstrate any protection against tau pathology or cognitive deficits. These findings indicate that different pathological tau epitopes, when used as the immunogen in vaccine strategies, confer differing levels of protection against tau pathology and disease progression in the rTg4510 animal model of tauopathy. This highlights the need for characterization of numerous pTau-therapeutic targets to identify the strongest candidates, or combinations thereof, for the development of successful immunotherapeutics against tauopathies.

## Results

### Qβ-AT8 and Qβ-PHF1 vaccines generate robust antibody responses

The Qβ-AT8 and Qβ-PHF1 vaccines were produced by conjugating the respective pTau peptides to pre-formed Qβ VLPs as described previously.^29,55,56^ Briefly, synthetic pTau peptides containing either the AT8 (pS199/pS202) or PHF1 (pS396/pS404) phosphorylation sites, with 6-7 amino acid residues flanking each phosphorylation site, and an added Gly-Gly-Cys spacer sequence (Fig 1a) were conjugated to surface-exposed lysine residues on the Qβ bacteriophage VLPs using a bifunctional cross-linker, succinimidyl 6-((beta-maleimidopropionamido) hexanoate) (SMPH) (Fig 1b). By assessing the mobility shift in bands of Qβ-conjugated pTau peptides compared to Qβ controls in a denaturing SDS-PAGE gel, we estimated ∼80 AT8 tau peptides or ∼150 PHF1 tau peptides are displayed per assembled Qβ-VLP which is composed of 180 monomers of the Qβ coat protein (Fig 1c). Thus, the Qβ-PHF1 VLP demonstrated a substantially higher conjugation efficiency than the Qβ-AT8 VLP.

**Fig 1.**
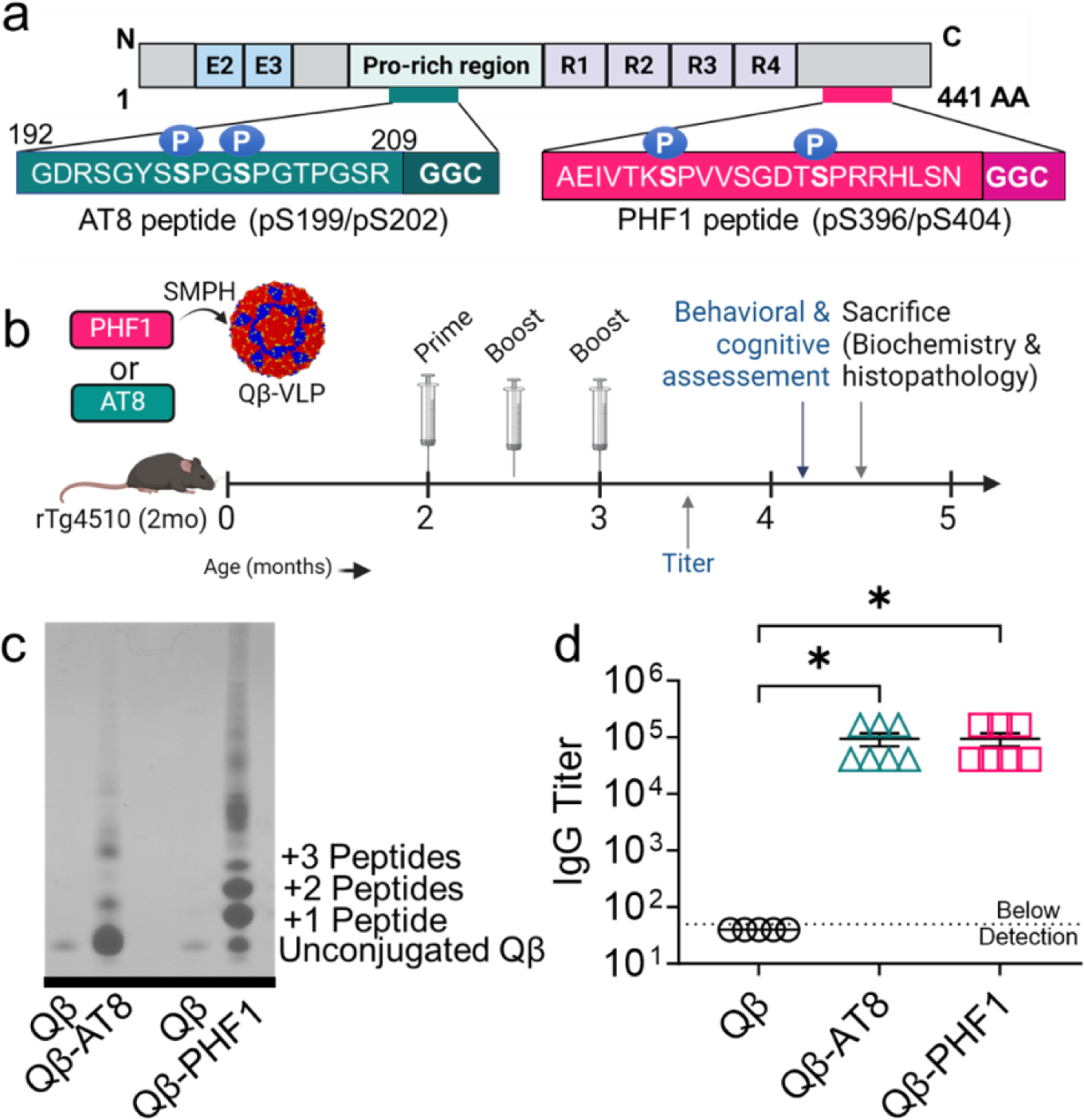
Qβ-AT8 and Qβ-PHF1 VLP conjugation, treatment timeline, and antibody titer responses. **a.** A 21-mer pTau peptide containing the Ser199/Ser202 phosphorylation sites and a modified Gly-Gly-Cys (GGC) C-terminal sequence (AT8 peptide) and a 25-mer pTau peptide containing the Ser396/Ser404 phosphorylation sites and a modified GGC C-terminal sequence (PHF1 peptide) were individually conjugated to surface exposed Lys residues of coat proteins on Qβ bacteriophage VLPs using SMPH cross-linker. **b.** 2-month-old rTg4510 mice received three bi-weekly intramuscular vaccinations with either unconjugated Qβ Control VLP (n=5), Qβ-AT8 VLP (n=7), or Qβ-PHF1 VLP (n=8) and were evaluated at 4.5-months for vaccine efficiency. **c.** An upward mobility shift on a 10% SDS denaturing gel shows the number of AT8 or PHF1 peptides conjugated per Qβ coat protein monomer; Qβ-PHF1 VLP shows a higher conjugation efficiency than Qβ-AT8 VLPs. **d.** Qβ-AT8 and Qβ-PHF1 VLP vaccines elicited significantly elevated serum IgG antibody titers at 6-weeks post-vaccination compared to Qβ Control. Graph displays mean ± SEM. p<0.05*; one-way ANOVA with Dunnett’s multiple comparisons. Qβ Control VLP (n=5), Qβ-AT8 VLP (n=7), or Qβ-PHF1 VLP (n=7).

To assess the immunogenicity and efficacy of the Qβ-AT8 and Qβ-PHF1 vaccines, 2-month-old transgenic rTg4510 tauopathy mice were intramuscularly immunized three-times with Qβ-AT8, Qβ-PHF1, or, as a control, unconjugated Qβ VLPs at biweekly intervals (Fig 1b). Two weeks after the final dose, serum IgG antibody titers against the respective phosphorylated tau peptides were assessed. Both the Qβ-AT8 and Qβ-PHF1 vaccines elicited high titer IgG responses against their respective tau peptide targets whereas the Qβ Control group had no detectable antibodies against phosphorylated tau (Fig 1d). Cognitive behavioral assessment was performed approximately one month later and then brain tissue was harvested for biochemical and neuropathological assessment (Fig 1b).

### Qβ-PHF1, but not Qβ-AT8, vaccination showed modest rescue of hippocampal-dependent memory

To assess the effects of Qβ-AT8 and Qβ-PHF1 vaccination on cognitive function in the rTg4510 tauopathy model, we conducted the Novel Object Recognition (NOR) and Morris Water Maze (MWM) behavioral assessments approximately one month after the final vaccine dose. Previous work^53,54,57^ demonstrated that behavioral impairments in rTg4510 mice develop over the 4-12-month period with a marked increase in hyperactivity occurring after 5-months potentially due to disruption of the *Fgf14* gene and six other genes by the bitransgenic insertion of the CaMK2a-tTA and tetO-MAPT*P301L transgenes.^58,59^ All behavioral assessments were performed as close to 4-months of age as possible to avoid potential confounding risks of hyperactivity.

We first conducted the NOR behavioral task to assess changes in short-term memory.^60^ On the first day, the animals are acclimated to the trial arena. 24-hours later (sample day), two identical objects are placed in the arena and the amount of time the animals spend investigating each object is recorded. 24-hours later (test day), one of the familiar objects is replaced with a novel object and the amount of time spent investigating each object is again recorded. On the sample day, all groups spent equal time investigating both objects indicating no preference for either object (Fig 2a). On the test day, as expected, non-transgenic healthy C57Bl6j (B6) mice spent a greater amount of time investigating the novel object, indicating intact recognition memory. There was no difference between time spent with the novel object and familiar object in the Qβ Control vaccinated rTg4510 tauopathy mice, indicating delay-dependent short-term working memory deficits (Fig 2b). Notably, this deficit was rescued in the Qβ-PHF1 vaccinated rTg4510 mice, which spent more time with the novel object (Fig 2b). However, Qβ-AT8 vaccination failed to rescue the recognition memory deficits, demonstrated by no preference for the novel object over the familiar object by Qβ-AT8 vaccinated mice (Fig 2b).

**Fig 2.**
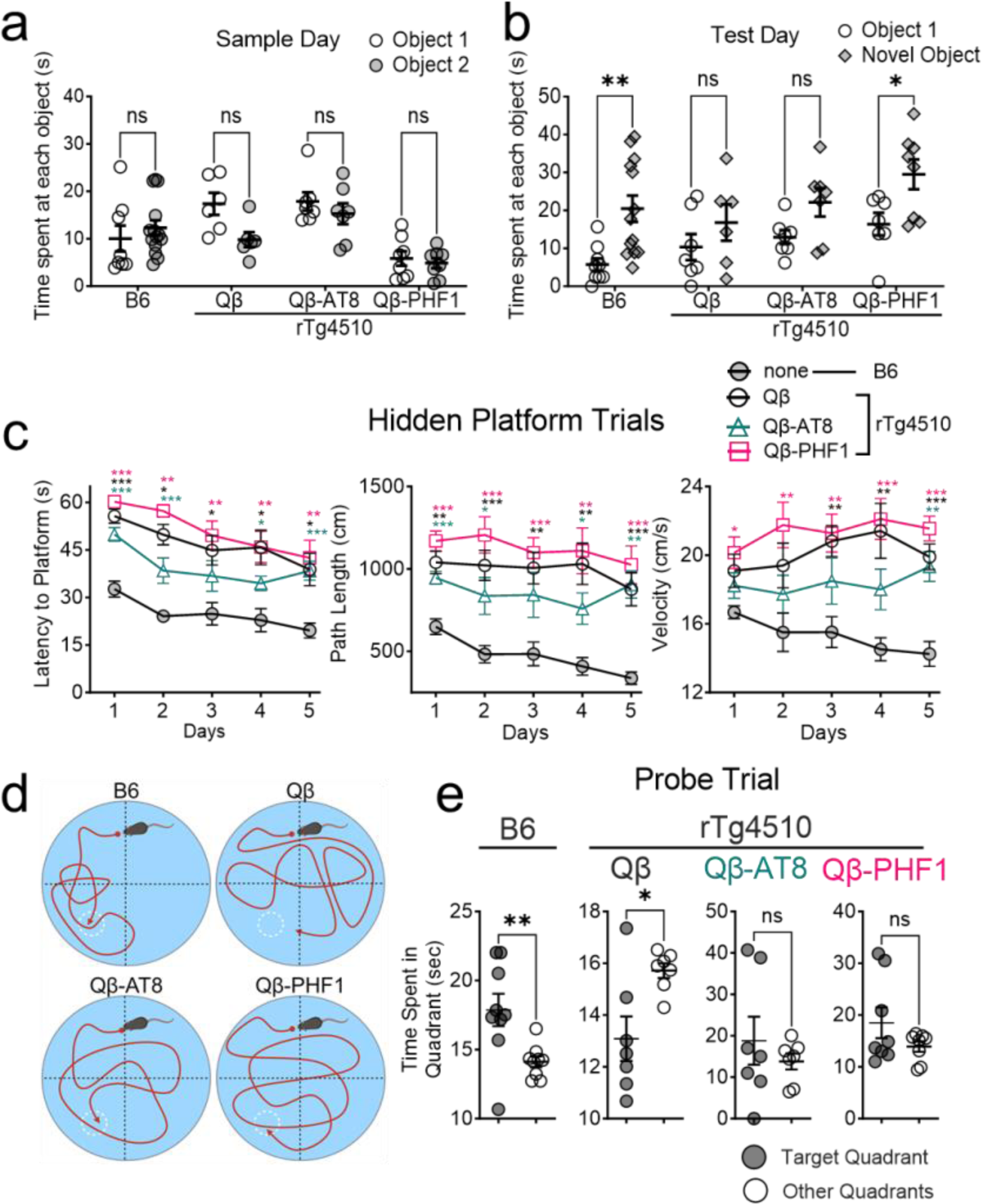
Qβ-PHF1 vaccination rescues delay-dependent memory deficits while Qβ-AT8 vaccination does not. **a.** In the Novel Object Recognition (NOR) test, no difference in the time spent with each object on the sample day. **b.** Twenty-four hours later (Test Day), non-transgenic (B6) mice spent significantly more time with the novel object. There was no difference in time spent with the novel object and familiar object for Qβ Control and Qβ-AT8 vaccinated rTg4510 mice, but this impairment was significantly rescued in Qβ-PHF1 vaccinated rTg4510 mice. **c.** Hidden platform trials of the Morris Water Maze (MWM) test show that B6 mice learned the location of the platform faster (shorter latency) than all rTg4510 vaccine groups. All rTg4510 vaccine groups exhibited hyperactivity based on distance traveled and velocity compared to B6 mice. **d-e.** During the probe trial (platform removed), B6 mice spent significantly more time exploring the target quadrant. Qβ Control rTg4510 mice spent significantly more time in the wrong quadrants. Qβ-AT8 and Qβ-PHF1 vaccinated rTg4510 spent equal time in all quadrants indicating a potentially milder impairment than Qβ Control mice but incomplete rescue of spatial memory. All graphs display mean ± SEM. Two-way ANOVA with Šidák correction (**a**, **b**). Two-way ANOVA with Dunnett’s correction comparing each group to the B6 group on each day (**c**). Student’s t-test (**e**). p<0.05*, p<0.01**, p<0.001***. B6 (n=9), Qβ Control (n=7), Qβ-AT8 (n=7), Qβ-PHF1 (n=8).

Next, we conducted the MWM behavior task to assess hippocampal-dependent spatial working memory consolidation. Over a 5-day period, mice are trained to find a hidden platform submerged under opacified water using visual cues in the room. For all groups, the latency to finding the platform decreased over each day indicating that the animals successfully learned the task of finding the platform (Fig 2c). All of the rTg4510 groups regardless of vaccine treatment exhibited a significantly longer latency to the platform than the non-transgenic B6 group potentially indicative of impaired working memory consolidation (Fig 2c). Additionally, all of the rTg4510 groups exhibited a longer path length and swimming velocity by the final training day which could potentially reflect the hyperactivity phenotype of this transgenic model. 24-hours after the conclusion of the 5-day training period, the hidden platform was removed (probe trial) and the amount of time the animals spent exploring the target quadrant (where the platform was previously located compared to the other quadrants) was recorded (Fig 2d). As expected, the non-transgenic B6 mice spent significantly more time in the target quadrant compared to the other quadrants indicating intact spatial memory consolidation based on the spatial cues on the wall. Comparatively, the Qβ Control vaccinated rTg4510 mice spent significantly more time exploring the other quadrants on average than the target quadrant indicating failure to consolidate the location of the platform after the 5-day training period (Fig 2e). There was no significant difference between time spent in the target quadrant and the other quadrants in the Qβ-AT8 and Qβ-PHF1 vaccinated rTg4510 groups (Fig 2e) indicating that these vaccines failed to rescue spatial memory deficits, however, the deficit may potentially have been less severe compared to the Qβ Control vaccinated rTg4510 group. Overall, the Qβ-PHF1 vaccinated rTg4510 group exhibited some rescue of hippocampal-dependent memory deficits observed in the Qβ Control vaccinated rTg4510 group while the Qβ-AT8 vaccine offered no protection against tauopathy-induced memory deficits.

### Qβ-PHF1 vaccination reduces soluble pathological tau levels in the brain but Qβ-AT8 vaccination does not

To evaluate the effect of Qβ-AT8 and Qβ-PHF1 vaccination on tau pathology we performed Western blot analysis of hippocampal lysates and immunohistochemical (IHC) staining on whole brain sections from vaccinated 4.5-month-old rTg4510 mice (1.5 months after the last boost). Hyperphosphorylation and aggregation of tau as NFTs has been shown to occur as early as 2-4 months of age in rTg4510 mice.^53,54,57^ Compared to Qβ Control vaccinated rTg4510 mice, the Qβ-PHF1 vaccinated rTg4510 mice showed significantly reduced AT180^+^ tau in the detergent soluble fraction (Fig 3a-b) while total tau levels (Tau5) remained unchanged indicating targeted reduction of AT180^+^ pTau (Fig 3a-b). There was no significant change in tau phosphorylated at the AT8 or PHF1 sites (Fig 3a-b). To validate the observed reduction in AT180^+^ pathological tau, we performed IHC staining of whole brain sections from the Qβ-PHF1 and Qβ Control vaccinated mice using the AT180 pathological tau marker. Compared to the Qβ Control vaccinated rTg4510 mice, the Qβ-PHF1 vaccinated rTg4510 mice exhibited a dramatic reduction in AT180 tau pathology in both the CA1 region of the hippocampus and the cerebral cortex (Fig 3c). Notable changes in tau pathology include reductions in extracellular tau staining, less dense peri-nuclear tau inclusions within the neurons, and a reduction in dystrophic neurite neuropathology in favor of organized dendrites in the hippocampus (Fig 3c). We also performed IHC staining in the Qβ-PHF1 and Qβ Control groups using the AT8 pTau marker but again found no differences (Supplemental Fig S1).

**Fig 3.**
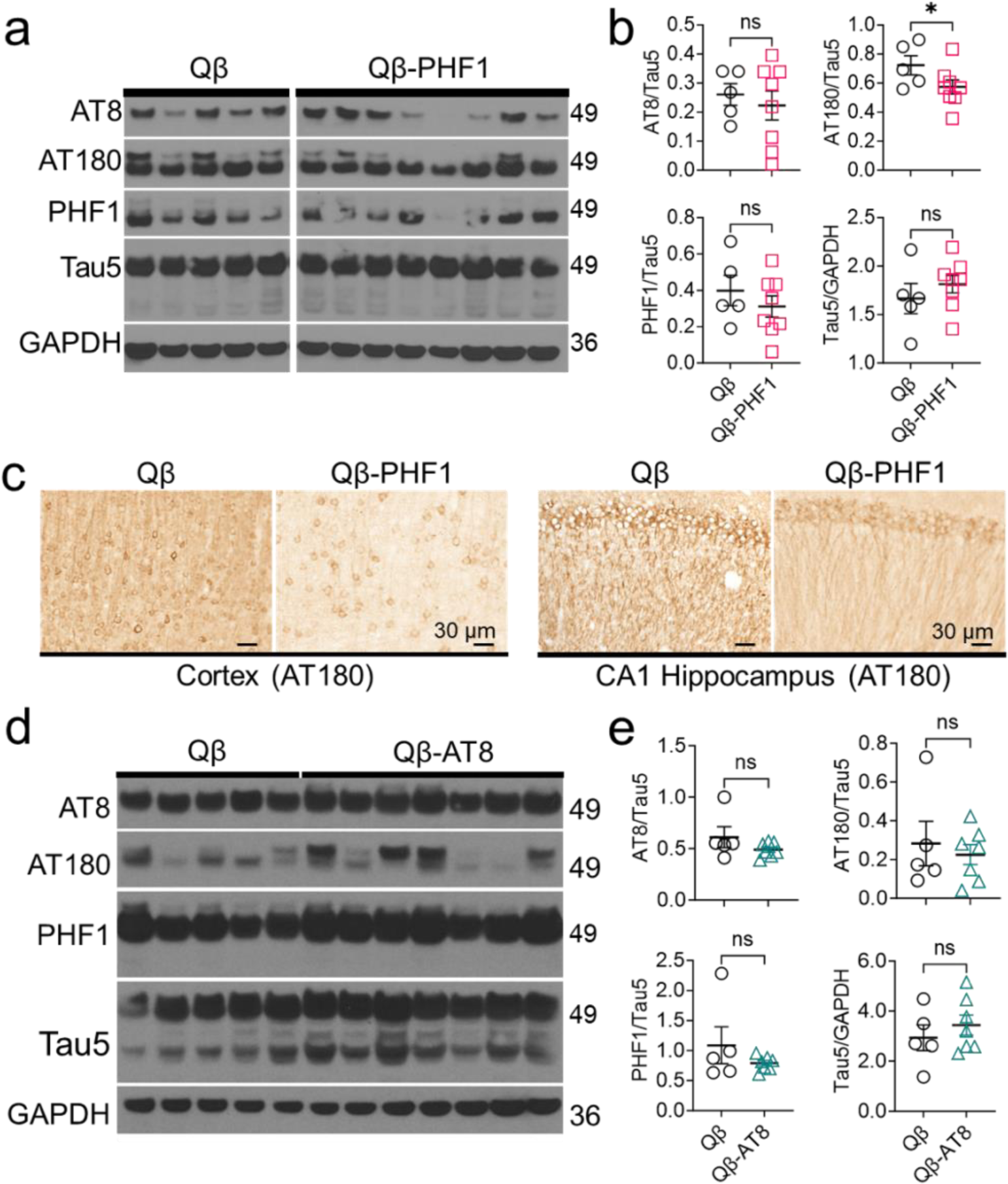
Qβ-PHF1 vaccination reduces soluble pathological tau in the brains of rTg4510 mice while Qβ-AT8 vaccination does not. **a.** Western blot of soluble hippocampal lysates from Qβ Control and Qβ-PHF1 vaccinated rTg4510 mice was evaluated for markers of phosphorylated (AT8, AT180, PHF1) and total tau (Tau5). All samples were run on the same gel. **b.** Compared to Qβ Control vaccinated mice, Qβ-PHF1 vaccinated mice showed significant reduction in pathological AT180 tau without any reduction in total physiologic tau. **c.** Immunohistochemistry of brain sections from Qβ Control and Qβ-PHF1 vaccinated rTg4510 mice was performed to validate the changes observed in AT180 tau where an obvious reduction in AT180 neuropathology was observed in both the cerebral cortex and CA1 hippocampus of Qβ-PHF1 vaccinated mice compared to the Qβ Control group. **d-e.** Western blot of soluble hippocampal lysates from Qβ Control and Qβ-AT8 vaccinated rTg4510 mice showed no reductions in pathological tau. All graphs display mean ± SEM. Student’s t-test (**b**, **e**). p<0.05*. Qβ Control (n=5), Qβ-AT8 (n=7), Qβ-PHF1 (n=8).

Western blot analysis of hippocampal lysates for the Qβ-AT8 vaccinated groups revealed no changes in soluble pathological tau levels compared to the Qβ Control group. There were no significant differences in AT8, AT180, or PHF1-positive pathological tau nor changes in total tau levels assessed by Tau5 indicating that Qβ-AT8 vaccination failed to reduce tau pathology in the rTg4510 model of tauopathy but also did not alter total tau levels (Fig 3d-e). Given our observations that the Qβ-AT8 vaccine failed to rescue cognitive deficits and similarly failed to reduce pathological tau levels, we decided not to pursue further investigation or characterization of this vaccine.

### Qβ-PHF1 vaccination preferentially reduces levels of insoluble tau aggregates in rTg4510 mice

To assess the effects of Qβ-PHF1 vaccination on insoluble tau levels we performed a Sarkosyl insoluble extraction of tau from the insoluble pellet fraction of the detergent-insoluble hippocampal lysates. Sarkosyl insoluble tau is composed of tau aggregates, the primary component of NFTs, which are correlated with disease progression and cognitive impairment.^35,61,62^ Using Western blot analysis, we evaluated the levels of phosphorylated insoluble tau (AT8) and total insoluble human tau (Tau12) compared to the levels of their non-aggregated Sarkosyl soluble counterparts. We found that the ratio of Sarkosyl insoluble to Sarkosyl soluble AT8 and Tau12-positive tau were both significantly reduced in the Qβ-PHF1 vaccinated group compared to the Qβ Control vaccinated group (Fig 4a-b). This data indicates that there is a preferential reduction in aggregated insoluble tau levels by Qβ-PHF1 vaccination.

**Fig 4.**
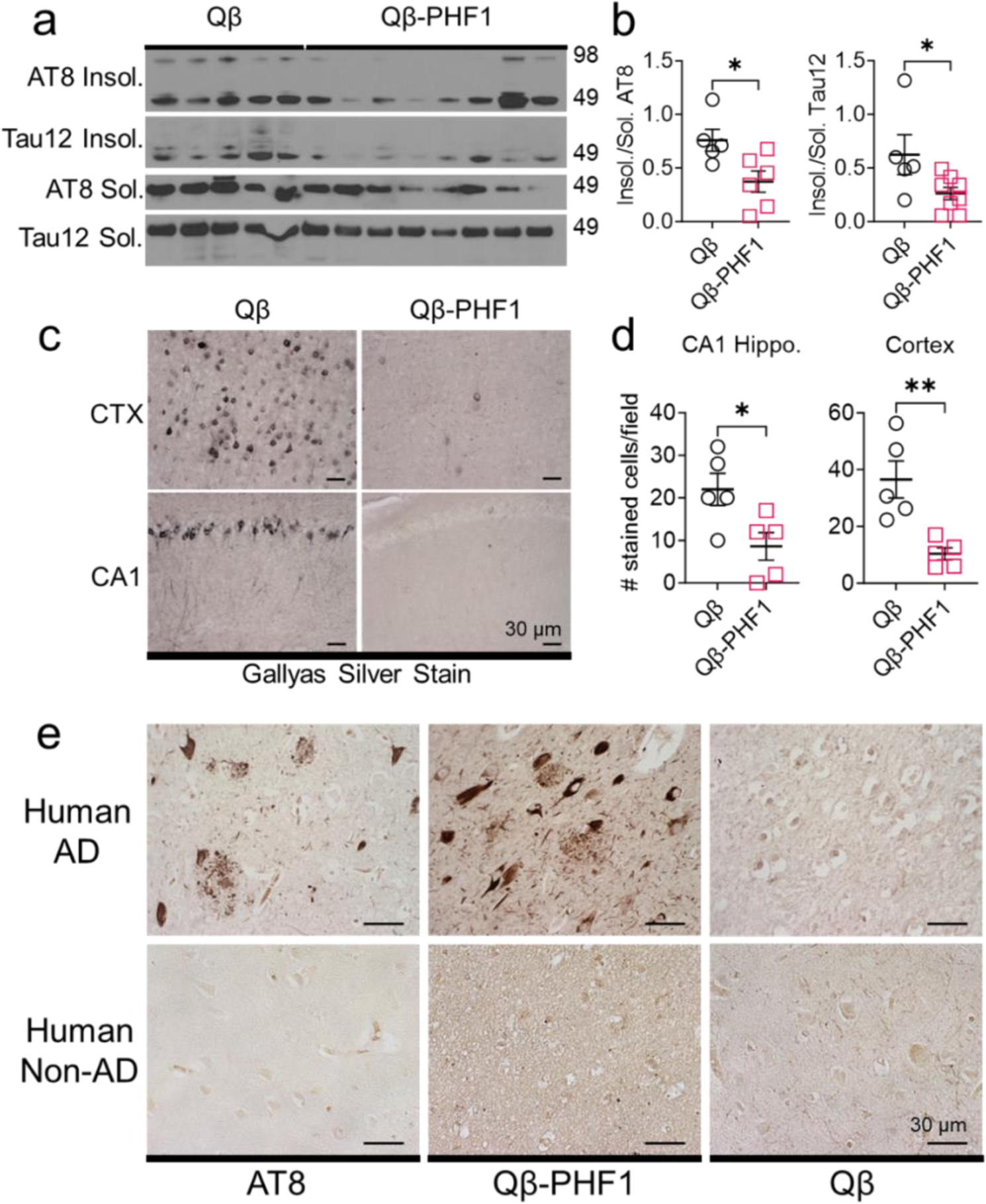
Qβ-PHF1 vaccination preferentially reduces Sarkosyl insoluble tau in rTg4510 mice and elicits antibodies specific to human pathological tau. **a-b**. Western blot (a) and quantification (b) showing a significant reduction in the ratio of Sarkosyl insoluble to soluble AT8 and Tau12 positive tau in the hippocampus of Qβ-PHF1 vaccinated rTg4510 mice compared to Qβ Control vaccinated mice. **c-d.** Gallyas silver impregnation (c) and quantification (d) showing reduction in insoluble tau and dramatic clearance of NFTs from the cerebral cortex and CA1 hippocampus of Qβ-PHF1 vaccinated mice compared to the Qβ Control group. **e.** Qβ-PHF1 immune sera stained somatodendritic NFTs, pre-tangles, neuritic plaques, neuropil threads, and ghost tangles in human autopsy AD, but not non-AD, brain tissue. Qβ Control sera did not show any specific staining. AT8 antibody staining was used as positive control. All graphs show mean ± SEM. Student’s t-test (**b**, **d**). p<0.05*, p<0.01**. **b.** Qβ Control (n=5), Qβ-PHF1 (n=8). **d.** Qβ Control (n=5), Qβ-PHF1 (n=5).

To further validate this finding, we performed Gallyas silver impregnation of whole brain sections from the Qβ-PHF1 and Qβ Control vaccinated rTg4510 mice to evaluate changes in NFT burden. Gallyas silver impregnation is the most sensitive and highly reproducible method for specifically staining NFTs whereby silver particles deposit on the argyrophilic NFT surface for histopathological visualization.^63,64^ In the 4.5-month-old Qβ Control vaccinated rTg4510 mice, there was an abundance of neurons staining positively for NFTs in both the CA1 region of the hippocampus and the cerebral cortex. Qβ-PHF1 vaccinated mice showed a dramatic reduction in the number of neurons positively stained using the Gallyas silver impregnation method in both the CA1 hippocampus and the cerebral cortex (Fig 4c-d). Taken together, the preferential reduction in Sarkosyl insoluble tau and NFT histopathology suggests that Qβ-PHF1 vaccination either preferentially promotes the clearance of NFTs from the brain or prevents the tau seeding and aggregation process that leads to the development NFTs within the brain.

### Qβ-PHF1 vaccination elicits antibodies reactive to pathological tau on human autopsy brain sections

To further demonstrate the specificity of antibody responses induced by Qβ-PHF1 vaccination for pathological tau, we stained human AD and non-demented human healthy control brain tissue sections with immune sera from the Qβ-PHF1 and Qβ Control vaccinated mice. We used the AT8 antibody as a positive control to validate the presence of tau pathology in the human AD brain and the lack of tau pathology in the human non-AD brain (Fig 4e). The human AD brain demonstrated numerous neuronal tau inclusions including pre-tangles, NFTs, and ghost tangles and dystrophic neurites including neuropil threads and neuritic plaques in concordance with the patient’s post-mortem pathological Braak staging of Braak VI. When stained using the whole immune sera from Qβ-PHF1 vaccinated mice, we observed robust staining of neuronal tau pathology including pre-tangles, NFTs, ghost tangles, neuropil threads, and neuritic plaques (Fig 4e). The immune sera from Qβ Control vaccinated mice did not detect any tau pathology in the human AD brain section (Fig 4e). The healthy human non-AD brain did not possess any tau inclusions as verified by AT8 staining. Importantly, no staining pattern was observed in the non-AD brain when stained with the Qβ-PHF1 vaccinated immune sera or with the Qβ Control immune sera (Fig 4e). This data further demonstrates that Qβ-PHF1 vaccination does not engage healthy physiological tau but instead preferentially engages pathological tau, with a propensity for aggregated tau suggestive of a conformation specific response.

We also performed staining of human AD hippocampal brain tissue sections with immune sera from Qβ-AT8 vaccinated mice which demonstrated specific detection of mature tau NFT pathology (Supplemental Fig S2). Notably, Qβ-AT8 immune sera only detected somatodendritic NFT and ghost tangle pathology but did not exhibit substantial staining of neuropil threads, neuritic plaques, or immature pre-tangle tau accumulations observed with the AT8 antibody.

### Qβ-PHF1 vaccination reduces reactive microgliosis in the rTg4510 model

Pathological tau has been shown to trigger a reactive inflammatory response in microglia and numerous genome-wide association studies have implicated maladaptive innate immune responses as underlying mechanisms in the neurodegenerative process of AD and related tauopathies.^65–73^ To evaluate the effect of Qβ-PHF1 vaccination on microglial histology in rTg4510 mice we performed IHC staining of whole brain slices using the microglial marker, Iba1, and the marker of microglial activation, CD45.^74^ We found that Qβ Control vaccinated rTg4510 mice exhibited a profound reactive microgliosis in both the CA1 region of the hippocampus and the cerebral cortex demonstrating a greater number of amoeboid microglia with increased positivity for CD45 (Fig 5a-b). Qβ-PHF1 vaccination showed a reduction in amoeboid microglia and concomitant increase in ramified microglia more characteristic of a homeostatic phenotype (Fig 5a). Additionally, Qβ-PHF1 vaccination resulted in a general reduction in CD45 staining relative to the Qβ Control group further suggestive of a reduction in tau-mediated reactive microgliosis (Fig 5b). This data suggests a reduction in microglial-mediated neuroinflammation as a consequence of pathological tau reduction by Qβ-PHF1 vaccination. Given that microglial dysfunction, including chronic inflammatory signaling, have been shown to contribute to further enhancement of tau accumulation in AD and tauopathy,^67,73,75–81^ this reduction in reactive microgliosis may be another mechanism by which Qβ-PHF1 vaccination protects against disease progression.

**Fig 5.**
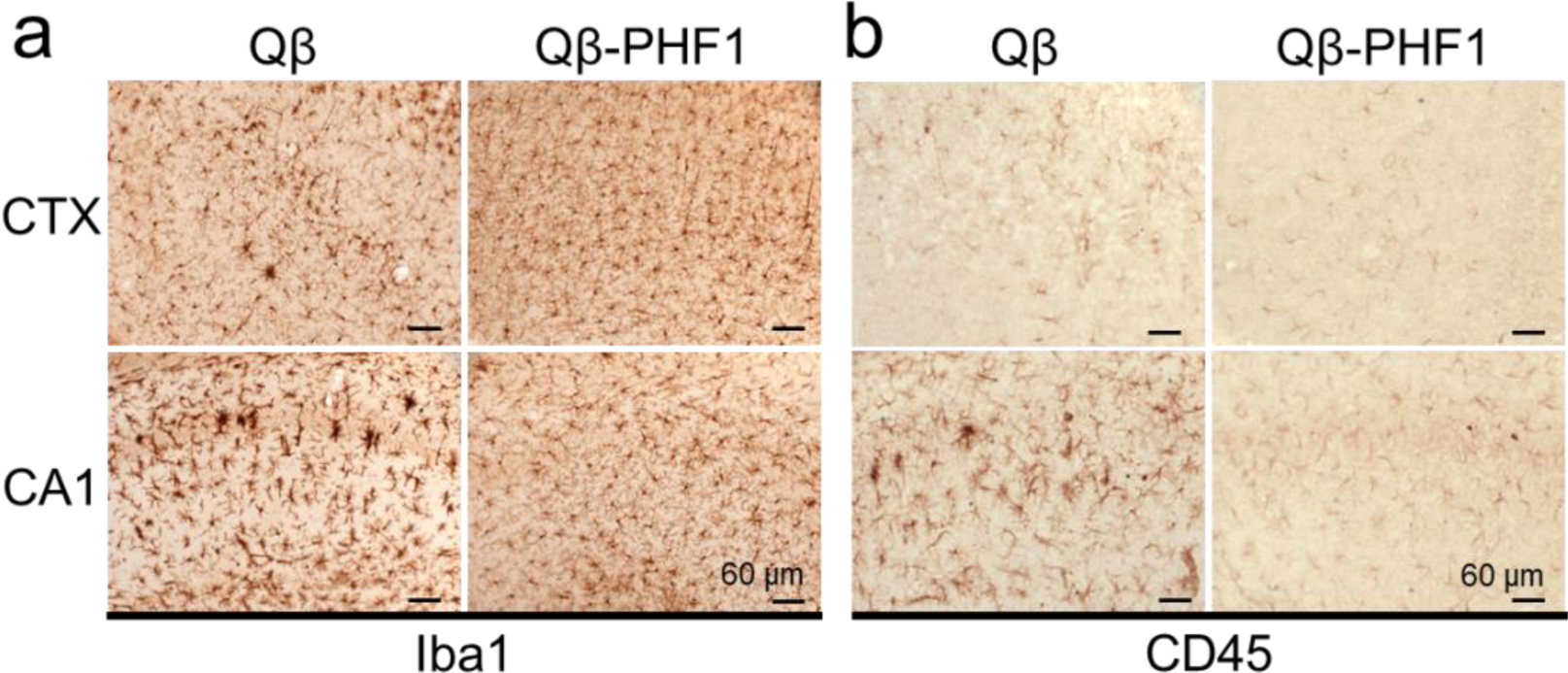
Qβ-PHF1 vaccination reduces reactive microgliosis in rTg4510 mice. **a**. IHC of Qβ Control and Qβ-PHF1 vaccinated rTg4510 mouse brain sections using Iba1 showed a reduction in amoeboid microglial morphology and overall Iba1 intensity in both the cerebral cortex and CA1 hippocampus of Qβ-PHF1 vaccinated mice compared to the Qβ Control group. **b**. Qβ-PHF1 vaccinated mice also exhibited reductions in CD45 intensity, a marker of microglial activation, in both the cerebral cortex and CA1 hippocampus compared to the Qβ Control group.

## Discussion

The central purpose of this investigation was to determine if either the AT8 or PHF1 phospho-tau epitopes, when displayed on Qβ VLPs, would be efficacious in generating targeted antibody responses, reducing tau pathology, and rescuing cognitive deficits in the rTg4510 model of tauopathy. We previously developed a Qβ VLP displaying the pT181 phospho-tau epitope (Qβ-pT181) which demonstrated a robust efficacy at reducing tau pathology, rescuing cognitive deficits, reducing neuronal cell death, and attenuating microglial inflammation. This epitope was selected because it is one of the first sites to undergo phosphorylation early in AD pathogenesis.^33^ Following similar rationale, we produced vaccines targeting the AT8 site and PHF1 site due to their importance as disease-specific phosphorylation sites. While both vaccines were highly immunogenic, eliciting high-titer antibody responses against their respective target in the vaccinated animals, only the Qβ-PHF1 vaccine was able to reduce tau pathology and rescue cognitive deficits. Immune sera from both the Qβ-PHF1 and Qβ-AT8 vaccines robustly detected tau pathology in human AD post-mortem brain tissue. However, the Qβ-AT8 immune sera exhibited a more limited range in tau pathology recognition (Supplementary Fig S2). This may explain the lack of tau clearance and cognitive protection by Qβ-AT8 in the rTg4510 mice compared to Qβ-PHF1.

Presently, we can only speculate why the PHF1-targeted vaccine was more efficacious than the AT8-targeted vaccine. Spatiotemporal differences in the AT8 and PHF1 phosphorylation sites may be a potential explanation. The AT8 and PHF1 sites are located on spatially distinct regions of the tau protein, within the more central proline-rich region (PRR) and the C-terminal region (CTR) respectively.^42,43^ As the tau protein oligomerizes and eventually forms large aggregate structures, it can adopt numerous conformations where these epitopes may be more or less accessible.^29^ Though the microtubule binding repeat domain (MTBR) is understood to be more involved in the amyloid core of paired helical filament tau structures leaving the PRR and CTR generally more accessible to antibody binding,^82,83^ better characterization of tau aggregate conformers will help elucidate whether these epitopes become buried or inaccessible during the aggregation process. It is worth noting that our previous Qβ-pT181 vaccine also targeted the PRR of tau. While the Qβ-AT8 vaccine failed to reduce tau pathology or rescue any cognitive deficits, the Qβ-pT181 vaccine was highly potent at reducing tau pathology and rescuing cognitive function, with enhanced efficacy compared to Qβ-PHF1.^29^

Temporal differences in the AT8 and PHF1 phosphorylation sites may better explain differences in vaccine efficacy. It has been well demonstrated that tau phosphorylation typically occurs in a sequential, site-specific manner during the disease process with some phosphorylation sites appearing early-on followed by additional PTMs as the disease progresses.^33,36,45^ While the AT8 antibody is now generally known to recognize the S202/T205 site,^42^ previous work identified S199/S202 site’s involvement in AT8 antibody recognition,^40,42,84^ some studies suggested S199/S202 phosphorylation may precede T205,^33^ and others identified S199/S202 as being a recognition site for Fyn interactions involved in tau pathogenesis.^85^ These are some of the reasons why we chose to target the S199/S202 site in this investigation. Recent findings, however, suggest that pS199 and pS202 are not elevated in AD brains relative to healthy controls until much later stages of disease, such as Braak stage V/VI.^45,46^ These pathologically modified epitopes may appear too late in the disease process for successful therapeutic targeting which could explain the failure of our Qβ-AT8 vaccine to reduce tau pathology and rescue cognition. Alternatively, the rapid disease progression and short timeline of our study in the rTg4510 model may have limited our ability to assess the efficacy of the Qβ-AT8 vaccine, because the pS199/pS202 epitopes may not have been abundantly present for therapeutic targeting. In contrast, the pS396/pS404 epitope has been shown to appear at somewhat earlier disease stages than the pS199/pS202 epitope, though it is also more abundant at later Braak stages.^33,36–38,44,45^ This difference potentially explains why the Qβ-PHF1 outperformed the Qβ-AT8 vaccine. Similarly, the pT181 tau epitope becomes pathologically modified earlier in the disease than both the AT8 and PHF1 sites which could explain why our previous Qβ-pT181 vaccine demonstrated the highest efficacy in the same animal model of tauopathy using the same vaccine dosing schedule. Taken together, our data potentially indicate that therapeutic targeting of earlier pathological tau epitopes in the disease process may yield the greatest protection against tau pathology and cognitive decline.

Lastly, differences in the pTau peptide conjugation efficiency on the Qβ VLPs may also explain the differences in efficacy between the Qβ-AT8 and Qβ-PHF1 vaccines. The Qβ-PHF1 vaccine demonstrated a substantially higher pTau peptide conjugation efficiency than the Qβ-AT8 vaccine (∼150 peptides/VLP compared to ∼80 peptides/VLP). The multivalent, repetitive antigenic display on VLPs is a major contributor to their immunogenicity.^31^ While both vaccines elicited the same titer of antibody responses against their pTau epitope, we did not assess if there are differences in the avidity of the antibody responses for their disease target. Interestingly, we speculate that the repetitive display of the PHF1 pTau peptides on the VLP may be mimicking a conformational epitope that is found within aggregated tau. This could potentially explain why the Qβ-PHF1 vaccine preferentially reduced insoluble tau aggregates as demonstrated by Sarkosyl-insoluble preparation of hippocampal lysates and silver impregnation of brain sections. Further characterization of the antibody responses to pTau VLP vaccination is necessary to better understand the specific binding interfaces on tau that are most beneficial for therapeutic targeting and to maximize antibody avidity for these interfaces. Future research could also employ cryo-electron microscopy techniques to better characterize the conformation of pTau peptides displayed on the VLP surface.

Additional characterization of the Qβ-AT8 and Qβ-PHF1 vaccines in other animal models of AD and tauopathy may be necessary to better understand the clinical translatability and clinical utility of these vaccines. The rTg4510 model is a highly aggressive tauopathy model which rapidly develops severe tau pathology. Alternative transgenic and knock-in human tauopathy animal models may provide more nuanced understanding of the efficacy of these vaccines in human AD and tauopathy. It is notable that the Qβ-PHF1 vaccine was able to demonstrate cognitive rescue and reduction of tau pathology in the rTg4510 model given the severity of pathology in this model. VLP vaccines targeting tau pathology show strong potential for a multi-epitope immunotherapy approach, by combining VLPs targeting multiple pTau epitopes, for addressing tau pathology in AD and primary tauopathies lending a sense of optimism that a future without these diseases is possible.

## Methods

### Production of Qβ-VLPs displaying phosphorylated S199/S202 (AT8) or phosphorylated S396/S404 (PHF1) tau peptides

Qβ VLPs were produced by purifying recombinantly expressed Qβ protein produced in *Escherichia coli* as described previously.^29,55,86^ Phosphorylated tau peptides were synthesized (American Peptide, USA) using sequences from the full-length (441 amino acid) isoform of human microtubule associated protein tau and modified with a glycine-glycine-cysteine C-terminal sequence to facilitate conjugation to the Qβ VLPs (AT8 sequence: GDRSGYS**pS**PG**pS**PGTPGSR-GGC; PHF1 sequence: AEIVTK**pS**PVVSGDT**pS**PRRHLSN-GGC) (Fig. 1a). SMPH (Thermo Fisher Scientific, catalog # 22363) was used as a bifunctional cross-linker to conjugate the terminal cysteine residue on each pTau peptide to the surface exposed lysine residues on the Qβ VLPs.^30,56^ Efficiency of conjugation was confirmed via mobility-shift gel electrophoresis on a 10% SDS denaturing polyacrylamide gel stained with Coomassie blue.

### Animal models and vaccine treatment paradigm

All animal work in this study was reviewed and approved by the University of New Mexico Institutional Animal Care and Use Committee (IACUC) under the following protocols (IACUC protocol #s: 19-200841-B-HSC (Breeding); 18-200761-HSC (Experimental)). Bi-transgenic rTg4510 were created by crossing the tet-transactivator line, Tg(Camk2a-tTA)1Mmay (JAX, stock# 007004), to the Tet responsive element line, Tg(tet0-MAPT*P301L)#Kha/J (JAX, stock# 015815).^53,57^ All transgenic rTg4510 and non-transgenic C57Bl/6j mice were also obtained from the Jackson Laboratory. Animals were housed in a specific pathogen free (SPF) facility, in a 12 h light/dark cycle with *ad libitum* access to food and water, in 85 in^2^ ventilated microisolator cages, supplemented with sterilized and autoclaved TEK fresh standard crinkle bedding; environmental enrichment included tissue paper and an elevated penthouse insert. Mice used in this study were not used for breeding and were housed by sex at a density of two to five mice per cage.

All animal work in this study was performed simultaneously with our previously published investigation of the Qβ-pT181 vaccine.^29^ As such, all animal behavior data from the B6 and Qβ groups (Fig 2), Qβ antibody titers (Fig 1d), and Qβ Control Western blot samples (Fig 3a) have been previously published,^29^ and were used here as matched controls in this study. The re-use of this data was permitted by the Creative Commons Attribution 4.0 International License (https://creativecommons.org/licenses/by/4.0/) and none of the data was changed from the original publication. All other data from the Qβ-AT8 and Qβ-PHF1 vaccine groups were generated exclusively for this study and have not been published previously. All animals chosen for experimental manipulation were healthy, of average weight, and had no history of rectal prolapse, skin dermatitis, or malocclusion. Beginning at 2-months, rTg4510 mice were treated with three, bi-weekly intramuscular injections into the rear hind-limb of either unconjugated Qβ (vaccine control) or Qβ-AT8 or Qβ-PHF1 (Fig 1) at an approximate concentration of 5 µg of VLP/injection. No adverse events were observed in any of the treatment groups.

Antibody titers were assessed two-weeks after the final injection. A battery of cognitive tests (NOR and MWM) were performed approximately one-month following the final vaccination. Following the conclusion of the cognitive testing, the animals were sacrificed by transcardial perfusion with ice-cold phosphate buffer under deep anesthesia with intraperitoneal Avertin injection. The perfused brains were immediately removed, the left hemispheres were immersion fixed in 4% paraformaldehyde (PFA; Electron Microscopy Services, catalog # RT15713) for neuropathological analysis via immunohistochemistry. The hippocampi from the right hemispheres were micro-dissected, weighed, snap frozen in liquid nitrogen, and stored at -80 C until subsequent biochemical analyses.

### Assessment of humoral immune responses in the serum

To assess antibody titer responses to vaccination, blood was collected using a retro-orbital capillary collection method two-weeks after the final vaccine dose. The blood was allowed to clot on ice for 20-minutes then centrifuged for 15 minutes at 15,000 *g* to separate the immune sera from the whole blood. Anti-pTau specific IgG titer responses were determined by endpoint dilution ELISA for each group of mice against their respective pTau peptide (AT8 peptide and PHF1 peptide). Briefly, Immulon-2 plates (Thermo Scientific) were incubated with 500 ng streptavidin (Thermo Fisher Scientific, catalog #434301) in pH 7.4 phosphate-buffered saline (PBS) overnight (ON) at 4 C. Following washing, SMPH was added to the wells at 1 ug/well and incubated for 2 hrs at room temperature (RT). In each group of experiments, each pTau peptide was added to the wells at 1 µg/well and incubated overnight at 4 C. The plate was subsequently blocked with 0.5% milk in PBS for 2 hrs. Four-fold dilutions of sera were added to each well and incubated for 2.5 hrs. The wells were probed with horseradish peroxidase (HRP)-conjugated secondary antibody [goat anti-mouse-IgG (Jackson ImmunoResearch, catalog #115-005-003; 1:4000)] for 1 hr. The reaction was developed using 3,3’, 5,5’-tetramethylbenzidine (TMB) substrate (Thermo Fisher Scientific, catalog # 34028) and stopped using 1 % HCl. Reactivity of sera for the target antigen was determined by measuring optical density at 450 nm (OD_450_). Wells with twice the OD_450_ of the background were considered to be positive and the highest dilution with a positive value was considered the end-point dilution titer.

### Novel Object Recognition and Morris Water Maze Behavioral Tests

For all behavioral experiments, animals were randomized to a blinded cage label so that the experimenter was blinded to the genotype and treatment of the animal. The NOR test involved a three-day paradigm for assessment of delay-dependent memory formation.^60^ On the first day, the animals were acclimated to a 75 cm^2^ arena which they freely explored for 5-minutes. On the second day, the animals again free explored the arena and became familiarized with two identical objects (glass jars) for 5-minutes. On the final day, one of the familiar objects was replaced with a novel object (plastic water bottle) and the animals freely explored the arena for 5-minutes. The percentage of time spent investigating each object was recorded using Ethovision XT8 (Noldus, Netherlands) live animal tracking with data processing in Excel and Prism.

The MWM test involved a six-day paradigm for assessment of learning and short-term spatial memory consolidation.^87,88^ All animals were trained for 5 days (4 daily trials, approximately 25 min inter-trial interval) to find a hidden platform submerged beneath opacified water at 26°C using spatial cues on the wall. During the acquisition phase, the animals freely navigated the tank of opaque water for 1 minute, and over 5 days (four trials daily) learned to navigate to a submerged platform using visual cues on the wall (visual cues were 2’ x 4’ made with different black/white shapes). Animals were randomized to search for a platform in a different area of the MWM, so results were counter-balanced; no bias was present to one quadrant over others. Animals were singly housed and kept warm in cages between trials. On the sixth day, a probe trial was performed by removing the platform and the percentage of time spent in the target quadrant vs. the other quadrants was recorded as a measurement of working spatial memory. All behavioral data was collected using Ethovision XT8 software (Noldus, Netherlands), and then processed in Excel and GraphPad Prism^®^.

### Immunohistochemistry

4% PFA-fixed hemi-brains were sectioned into 30-µm thick sagittal sections using a cryo-microtome. Free-floating sagittal sections derived from multiple mouse brains per group were utilized for all of the immunohistochemical analyses. First, sections were blocked in normal goat serum (NGS; Thermo Fisher Scientific) in 1x PBS pH7.4 with 0.4% Triton X-100 (PBST) for 1h at room temperature and included with primary antibodies. Antibody dilutions for each immunostain were as follows: AT180 (Thermo Fisher Scientific, MN1040) at 1:500, AT8 (Thermo Fisher Scientific, MN1020) at 1:500, Iba1 (WAKO, 019-19741) at 1:500, and CD45 (BioRad, MCA43R) at 1:250 incubated overnight at 4°C. Respective secondary antibodies conjugated to biotin (1:250, Jackson Immunoresearch) were used. Sections were incubated with Avidin/Biotin enzyme Complex (ABC reagent, Vector Laboratories; for immunohistochemistry) reagent for 1 h at room temperature. Immunoreactive signals were developed by incubating sections in 3-3’-diaminobenzadine (DAB) reagent (Vector Laboratories). Sections were mounted on glass slides which were serially dehydrated in ethanol, cleared with xylenes and mounted with Permount™ (Fischer Scientific). Bright field images were acquired using an Olympus BX-51 microscope (Olympus America Inc.), equipped with an Optronics digital camera and Picture Frame image capture software (Optronics).

### Gallyas Silver Impregnation

Gallyas silver impregnation staining was performed on 30-µm free-floating sagittal brain sections using a modified method for free-floating tissue samples as previously described.^29,89^ 30-μm thick sections were washed with water then incubated in 5% Periodic acid for 5 min, and washed with water again. Next, the sections were incubated for 1 min with a solution of alkaline silver iodide (containing sodium hydroxide, potassium iodide, and silver nitrate). Then the tissues were incubated for 10 min in 0.5% glacial acetic acid then developed in developer solution for up to 5 min. Developer working solution was mixed at a ratio of 3 volumes of Solution II to 10 volumes of Solution I followed by addition of 7 volumes of Solution III: Stock Solution I: 5% sodium carbonate; Stock Solution II: 0.2% Sodium nitrate, 0.2% silver nitrate, 1% tungstosilicic acid; Stock Solution III: 0.2% ammonium nitrate, 0.2% silver nitrate, 1% tungstosilicic acid, 73% wt/vol formaldehyde (neat). After developing the tissues were rinsed with 0.5% glacial acetic acid, rinsed with water, incubated with 0.1% gold chloride for 5 min, rinsed again with water, incubated with 1% sodium thiosulfate for 5 min, and finally rinsed once more with water. The silver impregnated sections were then mounted to glass slides, dried, rinsed carefully with xylenes and then glass coverslips were affixed with Permount™ (Fischer Scientific).

### Quantitative Morphometry of Gallyas Silver Impregnation

Bright field images of silver impregnated tissue sections were acquired at 40x magnification using an Olympus BX-51 microscope (Olympus America Inc.), equipped with an Optronics digital camera and Picture Frame image capture software (Optronics). CA1 hippocampus and overlying cerebral cortex were imaged from n=5 mice per group. The number of cells stained positively by silver impregnation were manually counted (one field of view for CA1 hippocampus; average of three fields of view for cortex) for each brain section.

### Characterization of Antibody Specificity in Human Brain Tissue

Clinically and neuropathologically diagnosed human post-mortem autopsy brain sections were obtained from Drs. Elaine Bearer and Karen SantaCruz. All tissue samples were obtained with consent from the patient by the University of New Mexico Office of the Medical Investigator. Neuropathologically confirmed AD brain tissue samples came from an 84-year-old female patient with Braak stage VI tau pathology. Neuropathologically confirmed non-AD brain tissue came from a 55-year-old female patient. Immune sera collected from vaccinated animals two-weeks after their final vaccination as previously stated was pooled (n=5 animals per group) and diluted (1:500) in 5% NGS in PBST and used as a primary antibody to stain human AD and non-demented age-matched control human brain tissue sections. 8 µM thick formalin-fixed paraffin embedded human brain tissue sections were deparaffinized in xylenes, serially rehydrated and then boiled in 10 mM Sodium Citrate Buffer pH 6.0 with 0.05% Tween-20 at 95°C for 30 min. The sections were washed with PBST, blocked in 5% NGS in PBST, and incubated overnight at 4°C with primary antibody (either pooled Qβ Control or Qβ-PHF1 sera, or AT8) at a 1:500 dilution. Secondary antibody incubations and DAB immunoreactive signal development occurred as stated previously.

### Western Blotting

Hippocampal brain tissue was homogenized for 1 min in Tissue Protein Extraction Reagent (TPER^®^, Thermo Scientific, catalog # 78510) at 10% w/v with phosphatase and protease inhibitor cocktails (Sigma Aldrich, P5726 and P8340, respectively), and then sonicated on ice for 30 s. Soluble hippocampal lysates were centrifuged at 12,000 *g* for 30 minutes at 4°C to separate the soluble lysates from the insoluble pellet. Soluble protein samples were prepared with NuPAGE™ lithium dodecyl sulfate (LDS) and reducing agent (RA) (Thermo fisher Scientific) and boiled at 95°C for 15 min. Samples were resolved via NuPAGE™ 4-12% Bis-Tris Polyacrylamide gradient gels (Thermo Fisher Scientific) and immunoblotted overnight in transfer buffer.^29^ All blots were processed in parallel and derived from the same experiment. Primary antibody dilutions were as follows: AT8 (Thermo Fisher Scientific (MN1020) 1:10,000; PHF1 (a kind gift from the late Peter Davies) 1:10,000; AT180 (Thermo Fisher Scientific, MN1040) 1:5,000; Tau5 (Millipore, MAB-361) 1:20,000; Tau12 (Millipore, MAB2241) 1:20,000; GAPDH (Millipore, AB2302) 1:20,000.

### Sarkosyl-Insoluble Tau Preparation

The Sarkosyl-insoluble fraction of tau was isolated from hippocampal samples as described previously. ^61,75^ The insoluble pellet generated from centrifugation of the 10% detergent soluble hippocampal lysates processed for Western blotting was sonicated in 10% wt/vol of cold buffer H (10 mM Tris-HCl, 1 mM EGTA, 0.8 mM NaCl, 10% sucrose, pH 7.4) supplemented by 0.1 mM PMSF (Sigma-Aldrich), with phosphatase and protease inhibitor cocktails (Sigma Aldrich, P5726 and P8340, respectively). The insoluble lysate was then centrifuged at 34,000×*g* in a Beckman Ti TLA-120.2 rotor for 30 min at 4°C. The supernatant was adjusted to 1% (w/v) N-laurylsarcosine (Sigma-Aldrich) with 1% (v/v) 2-mercaptoethanol (Sigma-Aldrich) and incubated at 37 °C for 2 hr with agitation. The sample was then ultra-centrifugated at 100,000 RPM for 35 min at room temperature. The Sarkosyl-soluble supernatant was then collected and resuspended in 1× NuPAGE™ LDS Sample Buffer (Thermo Fisher Scientific). The Sarkosyl-insoluble pellet was washed several times in 1% Sarkosyl solution prepared in cold buffer H and then resuspended in 1x NuPAGE™ LDS Sample Buffer. The Sarkosyl-soluble and insoluble fractions were loaded separately onto NuPAGE™ 4-12% Bis-Tris Polyacrylamide Gels (Thermo Fisher Scientific) and immunoblotted as described in the Western Blotting section. The dilutions of primary and secondary antibodies were the same as previously stated.

### Statistical Analysis and Study Blinding

Animal numbers for all experiments were determined by Power analysis prior to study initiation. Statistical analyses were performed using GraphPad Prism (USA) software. Unless otherwise noted, all data are presented as mean ± SEM. A student’s *t*-test was employed when two groups were being analyzed. A one-way ANOVA, with Tukey’s post-hoc test for multiple comparisons, was used when investigating three or more groups. Two-way ANOVA with either Šidák or Dunnett’s correction for post-hoc multiple comparisons was used for analysis of the Novel Object Recognition and Morris Water Maze tests. All significance values are reported at an α=0.05. Rout’s outlier analysis with a false discovery rate of 1% was performed on all data sets to identify and remove any outliers in all statistical analyses; number of animals analyzed in each test is indicated in the figure legends.

All experiments were performed using a unique pathology numbering system (for necropsy studies) to avoid any subjective bias. Experimenters were blinded to the genotype and vaccine treatment performed for all experiments in this study. Genotype and treatment information was only made available after the completion of the analyses.

## Data Availability

The datasets used and/or analyzed in the current study are available from the corresponding author upon reasonable request.

## Acknowledgments

We would like to thank Drs. Elaine Bearer and Karen SantaCruz for sharing neuropathologically characterized human post-mortem autopsy brain sections. We would like to thank the University of New Mexico Animal Resource Facility staff for the excellent care of our animals. We would like to thank Dr. Carissa Milliken, Dr. Russ Morton and Dr. Jonathan Brigman for assistance with behavioral data acquisition and for sharing their behavioral equipment. We would like to thank Dr. Shanya Jiang for assistance with Sarkosyl extraction of tau. We would also like to thank Myranda Thompson for assistance with Gallyas Silver Impregnation and immunohistochemistry.

## Author Contributions

J.H., N.M., B.C., and K.B. designed the study and drafted the manuscript. J.P. and B.C. engineered and conjugated the Qβ-AT8 and Qβ-PHF1 VLPs. N.M. and J.P vaccinated the mice and performed the ELISA analyses. K.B. performed the Sarkosyl extraction assays. N.M. and J.H. performed Western blot analyses. J.H. and N.M. performed immunohistochemical analyses. J.H. and K.B. performed Gallyas silver impregnation. N.M. performed behavioral testing and analysis.

## Supplemental material

**Supplementary Fig S1.**
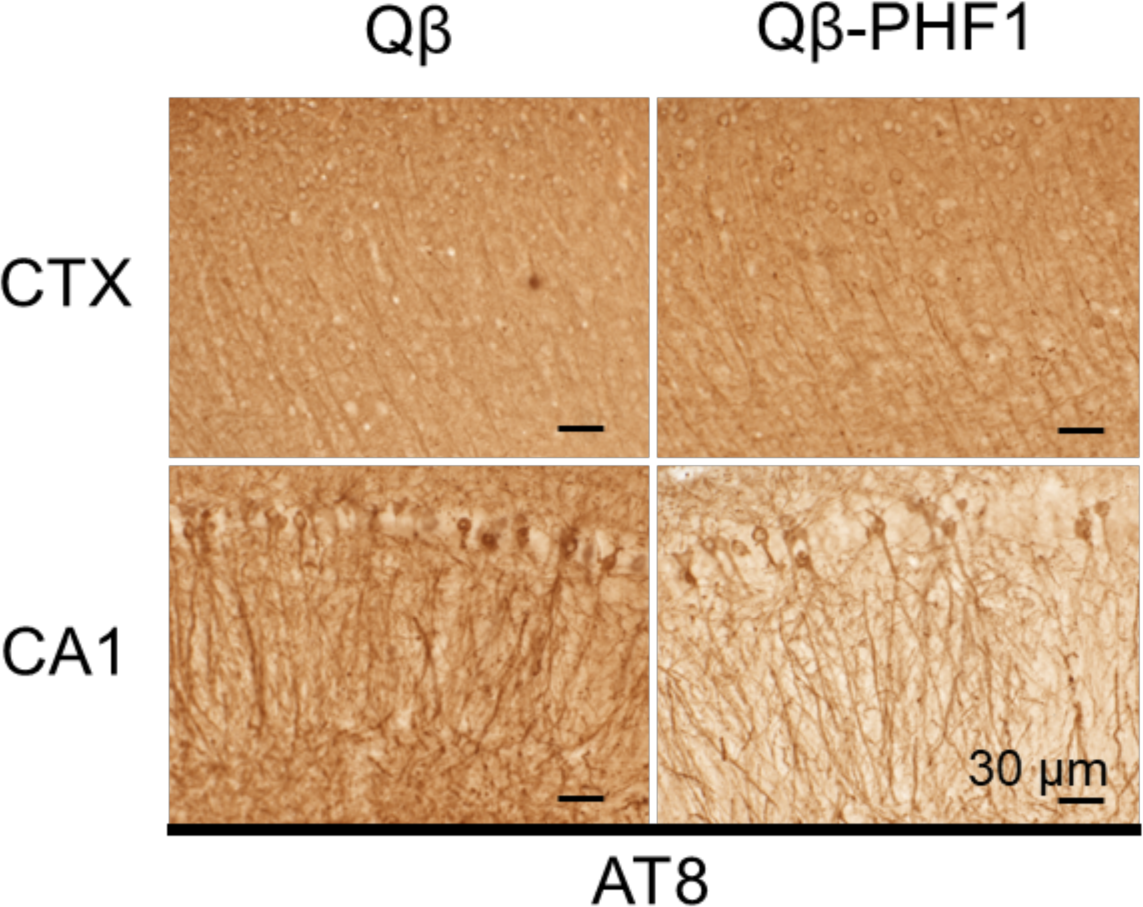
Qβ-PHF1 vaccination fails to reduce AT8 pathology in rTg4510 mice. IHC of brain sections from Qβ Control and Qβ-PHF1 vaccinated rTg4510 mice using AT8 was performed to validate the observations from Western blot analysis that soluble AT8 levels are unchanged in Qβ-PHF1 vaccinated mice compared to Qβ Controls. No obvious differences are observed in AT8 histopathology between Qβ Control and Qβ-PHF1 vaccinated rTg4510 mice in either the cerebral cortex or CA1 hippocampus.

**Supplementary Fig S2.**
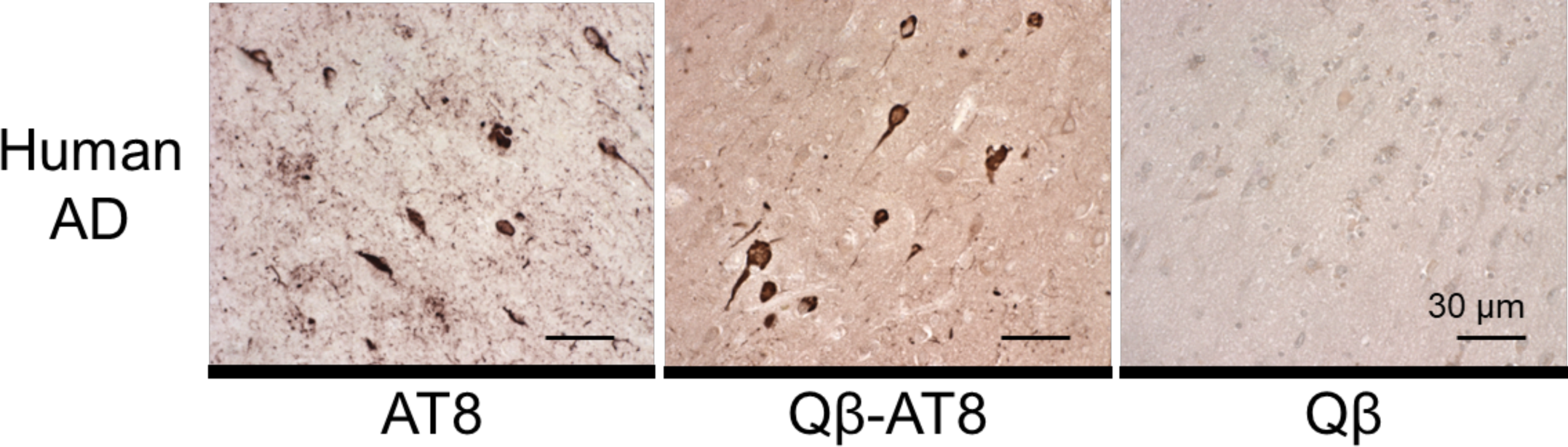
Immune sera from Qβ Control and Qβ-AT8 vaccinated mice was used to stain human post-mortem hippocampal sections from AD brain tissue and compared to AT8 histopathology. Qβ-AT8 immune sera stained somatodendritic NFT and ghost tangle histopathology in AD brain tissue similarly to AT8 but did not robustly stain other pathological features observed in AT8 such as neuropil threads or neuritic plaques. Qβ Control sera did not exhibit specific staining of any tau pathology.

## References

1. Alzheimer, A., Stelzmann, R. A., Schnitzlein, H. N. & Murtagh, F. R. An English translation of Alzheimer’s 1907 paper, ‘Uber eine eigenartige Erkankung der Hirnrinde’. Clin Anat 8, 429–431 (1995).

2. DeTure, M. A. & Dickson, D. W. The neuropathological diagnosis of Alzheimer’s disease. Molecular Neurodegeneration 14, 32 (2019).

3. Iqbal, K., Liu, F., Gong, C.-X. & Grundke-Iqbal, I. Tau in Alzheimer Disease and Related Tauopathies. Curr Alzheimer Res 7, 656–664 (2010).

4. Alonso, A. del C., Zaidi, T., Novak, M., Grundke-Iqbal, I. & Iqbal, K. Hyperphosphorylation induces self-assembly of τ into tangles of paired helical filaments/straight filaments. Proc Natl Acad Sci U S A 98, 6923–6928 (2001).

5. Pevalova, M., Filipcik, P., Novak, M., Avila, J. & Iqbal, K. Post-translational modifications of tau protein. Bratisl Lek Listy 107, 346–353 (2006).

6. Wilcock, G. K. & Esiri, M. M. Plaques, tangles and dementia. A quantitative study. J Neurol Sci 56, 343–356 (1982).

7. Arriagada, P. V., Growdon, J. H., Hedley-Whyte, E. T. & Hyman, B. T. Neurofibrillary tangles but not senile plaques parallel duration and severity of Alzheimer’s disease. Neurology 42, 631–639 (1992).

8. Ossenkoppele, R. et al. Amyloid and tau PET-positive cognitively unimpaired individuals are at high risk for future cognitive decline. Nat Med 28, 2381–2387 (2022).

9. Hall, B. et al. In vivo tau PET imaging in dementia: Pathophysiology, radiotracer quantification, and a systematic review of clinical findings. Ageing Res Rev 36, 50–63 (2017).

10. La Joie, R. et al. Prospective longitudinal atrophy in Alzheimer’s disease correlates with the intensity and topography of baseline tau-PET. Sci Transl Med 12, eaau5732 (2020).

11. Vogel, J. W. et al. Spread of pathological tau proteins through communicating neurons in human Alzheimer’s disease. Nat Commun 11, 2612 (2020).

12. Goedert, M., Eisenberg, D. S. & Crowther, R. A. Propagation of Tau Aggregates and Neurodegeneration. Annu Rev Neurosci 40, 189–210 (2017).

13. Frost, B. & Diamond, M. I. Prion-like mechanisms in neurodegenerative diseases. Nat Rev Neurosci 11, 155–159 (2010).

14. He, Z. et al. Amyloid-β plaques enhance Alzheimer’s brain tau-seeded pathologies by facilitating neuritic plaque tau aggregation. Nat Med 24, 29–38 (2018).

15. Liu, L. et al. Trans-Synaptic Spread of Tau Pathology In Vivo. PLOS ONE 7, e31302 (2012).

16. Iba, M. et al. Synthetic tau fibrils mediate transmission of neurofibrillary tangles in a transgenic mouse model of Alzheimer’s-like tauopathy. J Neurosci 33, 1024–1037 (2013).

17. Clavaguera, F. et al. Brain homogenates from human tauopathies induce tau inclusions in mouse brain. Proc Natl Acad Sci U S A 110, 9535–9540 (2013).

18. DeVos, S. L. et al. Synaptic Tau Seeding Precedes Tau Pathology in Human Alzheimer’s Disease Brain. Front. Neurosci. 12, (2018).

19. Rodrigues, S. et al. Spreading of Tau Protein Does Not Depend on Aggregation Propensity. J Mol Neurosci 73, 693–712 (2023).

20. van Dyck, C. H. et al. Lecanemab in Early Alzheimer’s Disease. N Engl J Med 388, 9–21 (2023).

21. Sims, J. R. et al. Donanemab in Early Symptomatic Alzheimer Disease: The TRAILBLAZER-ALZ 2 Randomized Clinical Trial. JAMA 330, 512–527 (2023).

22. Sifniotis, V., Cruz, E., Eroglu, B. & Kayser, V. Current Advancements in Addressing Key Challenges of Therapeutic Antibody Design, Manufacture, and Formulation. Antibodies (Basel) 8, 36 (2019).

23. Jönsson, L. et al. The affordability of lecanemab, an amyloid-targeting therapy for Alzheimer’s disease: an EADC-EC viewpoint. Lancet Reg Health Eur 29, 100657 (2023).

24. Cummings, J. et al. Lecanemab: Appropriate Use Recommendations. J Prev Alzheimers Dis 10, 362–377 (2023).

25. Mena Lora, A. J., et al. Feasibility and impact of a monoclonal antibody infusion program in reaching vulnerable underserved communities. Infect Control Hosp Epidemiol 44, 1690–1692.

26. Manly, J. J. & Deters, K. D. Donanemab for Alzheimer Disease—Who Benefits and Who Is Harmed? JAMA 330, 510–511 (2023).

27. Nooraei, S. et al. Virus-like particles: preparation, immunogenicity and their roles as nanovaccines and drug nanocarriers. Journal of Nanobiotechnology 19, 59 (2021).

28. Farlow, M. R. et al. Long-term treatment with active Aβ immunotherapy with CAD106 in mild Alzheimer’s disease. Alzheimers Res Ther 7, 23 (2015).

29. Maphis, N. M. et al. Qß Virus-like particle-based vaccine induces robust immunity and protects against tauopathy. NPJ Vaccines 4, 26 (2019).

30. Chackerian, B. Virus-like particles: flexible platforms for vaccine development. Expert Review of Vaccines 6, 381–390 (2007).

31. Bachmann, M. F. & Jennings, G. T. Vaccine delivery: a matter of size, geometry, kinetics and molecular patterns. Nat Rev Immunol 10, 787–796 (2010).

32. Jennings, G. T. & Bachmann, M. F. Immunodrugs: therapeutic VLP-based vaccines for chronic diseases. Annu Rev Pharmacol Toxicol 49, 303–326 (2009).

33. Wesseling, H. et al. Tau PTM Profiles Identify Patient Heterogeneity and Stages of Alzheimer’s Disease. Cell 183, 1699–1713.e13 (2020).

34. Braak, E., Braak, H. & Mandelkow, E. M. A sequence of cytoskeleton changes related to the formation of neurofibrillary tangles and neuropil threads. Acta Neuropathol 87, 554–567 (1994).

35. Braak, H., Thal, D. R., Ghebremedhin, E. & Del Tredici, K. Stages of the pathologic process in Alzheimer disease: age categories from 1 to 100 years. J Neuropathol Exp Neurol 70, 960–969 (2011).

36. Koss, D. J. et al. Soluble pre-fibrillar tau and β-amyloid species emerge in early human Alzheimer’s disease and track disease progression and cognitive decline. Acta Neuropathol 132, 875–895 (2016).

37. Mondragón-Rodríguez, S., Perry, G., Luna-Muñoz, J., Acevedo-Aquino, M. C. & Williams, S. Phosphorylation of tau protein at sites Ser(396-404) is one of the earliest events in Alzheimer’s disease and Down syndrome. Neuropathol Appl Neurobiol 40, 121–135 (2014).

38. Augustinack, J. C., Schneider, A., Mandelkow, E.-M. & Hyman, B. T. Specific tau phosphorylation sites correlate with severity of neuronal cytopathology in Alzheimer’s disease. Acta Neuropathol 103, 26–35 (2002).

39. Lasagna-Reeves, C. A. et al. Identification of oligomers at early stages of tau aggregation in Alzheimer’s disease. FASEB J 26, 1946–1959 (2012).

40. Porzig, R., Singer, D. & Hoffmann, R. Epitope mapping of mAbs AT8 and Tau5 directed against hyperphosphorylated regions of the human tau protein. Biochem Biophys Res Commun 358, 644–649 (2007).

41. Malia, T. J. et al. Epitope mapping and structural basis for the recognition of phosphorylated tau by the anti-tau antibody AT8. Proteins 84, 427–434 (2016).

42. Goedert, M., Jakes, R. & Vanmechelen, E. Monoclonal antibody AT8 recognises tau protein phosphorylated at both serine 202 and threonine 205. Neurosci Lett 189, 167–169 (1995).

43. Otvos, L. et al. Monoclonal antibody PHF-1 recognizes tau protein phosphorylated at serine residues 396 and 404. J Neurosci Res 39, 669–673 (1994).

44. Zhou, X.-W. et al. Assessments of the accumulation severities of amyloid β-protein and hyperphosphorylated tau in the medial temporal cortex of control and Alzheimer’s brains. Neurobiology of Disease 22, 657–668 (2006).

45. Neddens, J. et al. Phosphorylation of different tau sites during progression of Alzheimer’s disease. Acta Neuropathologica Communications 6, 52 (2018).

46. Lantero-Rodriguez, J. et al. CSF p-tau205: a biomarker of tau pathology in Alzheimer’s disease. Acta Neuropathol 147, 12 (2024).

47. Abraha, A. et al. C-terminal inhibition of tau assembly in vitro and in Alzheimer’s disease. J Cell Sci 113 **Pt 21**, 3737–3745 (2000).

48. Rosenqvist, N. et al. Highly specific and selective anti-pS396-tau antibody C10.2 targets seeding-competent tau. Alzheimers Dement (N Y) 4, 521–534 (2018).

49. Chukwu, J. E. et al. Tau Antibody Structure Reveals a Molecular Switch Defining a Pathological Conformation of the Tau Protein. Sci Rep 8, 6209 (2018).

50. Jackson, S. J. et al. Short Fibrils Constitute the Major Species of Seed-Competent Tau in the Brains of Mice Transgenic for Human P301S Tau. J Neurosci 36, 762–772 (2016).

51. Jeganathan, S. et al. Proline-directed pseudo-phosphorylation at AT8 and PHF1 epitopes induces a compaction of the paperclip folding of Tau and generates a pathological (MC-1) conformation. J Biol Chem 283, 32066–32076 (2008).

52. Braak, H., Alafuzoff, I., Arzberger, T., Kretzschmar, H. & Del Tredici, K. Staging of Alzheimer disease-associated neurofibrillary pathology using paraffin sections and immunocytochemistry. Acta Neuropathol 112, 389–404 (2006).

53. Ramsden, M. et al. Age-Dependent Neurofibrillary Tangle Formation, Neuron Loss, and Memory Impairment in a Mouse Model of Human Tauopathy (P301L). J. Neurosci. 25, 10637–10647 (2005).

54. SantaCruz, K. et al. Tau Suppression in a Neurodegenerative Mouse Model Improves Memory Function. Science 309, 476–481 (2005).

55. Crossey, E. et al. A cholesterol-lowering VLP vaccine that targets PCSK9. Vaccine 33, 5747–5755 (2015).

56. Chackerian, B., Rangel, M., Hunter, Z. & Peabody, D. S. Virus and virus-like particle-based immunogens for Alzheimer’s disease induce antibody responses against amyloid-beta without concomitant T cell responses. Vaccine 24, 6321–6331 (2006).

57. Spires, T. L. et al. Region-specific Dissociation of Neuronal Loss and Neurofibrillary Pathology in a Mouse Model of Tauopathy. The American Journal of Pathology 168, 1598–1607 (2006).

58. Goodwin, L. O. et al. Large-scale discovery of mouse transgenic integration sites reveals frequent structural variation and insertional mutagenesis. Genome Res 29, 494–505 (2019).

59. Gamache, J. et al. Factors other than hTau overexpression that contribute to tauopathy-like phenotype in rTg4510 mice. Nat Commun 10, 2479 (2019).

60. Antunes, M. & Biala, G. The novel object recognition memory: neurobiology, test procedure, and its modifications. Cogn Process 13, 93–110 (2012).

61. Greenberg, S. G. & Davies, P. A preparation of Alzheimer paired helical filaments that displays distinct tau proteins by polyacrylamide gel electrophoresis. Proceedings of the National Academy of Sciences 87, 5827–5831 (1990).

62. Kidd, M. Paired helical filaments in electron microscopy of Alzheimer’s disease. Nature 197, 192–193 (1963).

63. Braak, H., Braak, E., Ohm, T. & Bohl, J. Silver impregnation of Alzheimer’s neurofibrillary changes counterstained for basophilic material and lipofuscin pigment. Stain Technol 63, 197–200 (1988).

64. Uchihara, T. Silver diagnosis in neuropathology: principles, practice and revised interpretation. Acta Neuropathol 113, 483–499 (2007).

65. Udeochu, J. C. et al. Tau activation of microglial cGAS-IFN reduces MEF2C-mediated cognitive resilience. Nat Neurosci 26, 737–750 (2023).

66. Griciuc, A. & Tanzi, R. E. The role of innate immune genes in Alzheimer’s disease. Curr Opin Neurol 34, 228–236 (2021).

67. Maphis, N. et al. Reactive microglia drive tau pathology and contribute to the spreading of pathological tau in the brain. Brain 138, 1738–1755 (2015).

68. Efthymiou, A. G. & Goate, A. M. Late onset Alzheimer’s disease genetics implicates microglial pathways in disease risk. Molecular Neurodegeneration 12, 43 (2017).

69. Prater, K. E. et al. Human microglia show unique transcriptional changes in Alzheimer’s disease. Nat Aging 3, 894–907 (2023).

70. Kunkle, B. W. et al. Genetic meta-analysis of diagnosed Alzheimer’s disease identifies new risk loci and implicates Aβ, tau, immunity and lipid processing. Nat Genet 51, 414–430 (2019).

71. Sims, R., Hill, M. & Williams, J. The multiplex model of the genetics of Alzheimer’s disease. Nat Neurosci 23, 311–322 (2020).

72. Reed, E. G. & Keller-Norrell, P. R. Minding the Gap: Exploring Neuroinflammatory and Microglial Sex Differences in Alzheimer’s Disease. Int J Mol Sci 24, 17377 (2023).

73. Jiang, S. et al. Proteopathic tau primes and activates interleukin-1β via myeloid-cell-specific MyD88- and NLRP3-ASC-inflammasome pathway. Cell Rep 36, 109720 (2021).

74. Jurga, A. M., Paleczna, M. & Kuter, K. Z. Overview of General and Discriminating Markers of Differential Microglia Phenotypes. Front Cell Neurosci 14, 198 (2020).

75. Bhaskar, K. et al. Regulation of tau pathology by the microglial fractalkine receptor. Neuron 68, 19–31 (2010).

76. Maphis, N. et al. Loss of tau rescues inflammation-mediated neurodegeneration. Front. Neurosci. 9, (2015).

77. Hulse, J. & Bhaskar, K. Crosstalk Between the NLRP3 Inflammasome/ASC Speck and Amyloid Protein Aggregates Drives Disease Progression in Alzheimer’s and Parkinson’s Disease. Frontiers in Molecular Neuroscience 15, (2022).

78. Ising, C. et al. NLRP3 inflammasome activation drives tau pathology. Nature 575, 669–673 (2019).

79. Venegas, C. & Heneka, M. T. Inflammasome-mediated innate immunity in Alzheimer’s disease. FASEB J 33, 13075–13084 (2019).

80. Heneka, M. T. et al. NLRP3 is activated in Alzheimer’s disease and contributes to pathology in APP/PS1 mice. Nature 493, 674–678 (2013).

81. Wang, C. et al. Microglial NF-κB drives tau spreading and toxicity in a mouse model of tauopathy. Nat Commun 13, 1969 (2022).

82. Fitzpatrick, A. W. P. et al. Cryo-EM structures of tau filaments from Alzheimer’s disease. Nature 547, 185–190 (2017).

83. Falcon, B. et al. Tau filaments from multiple cases of sporadic and inherited Alzheimer’s disease adopt a common fold. Acta Neuropathol 136, 699–708 (2018).

84. Biernat, J. et al. The switch of tau protein to an Alzheimer-like state includes the phosphorylation of two serine-proline motifs upstream of the microtubule binding region. EMBO J 11, 1593–1597 (1992).

85. Bhaskar, K., Yen, S.-H. & Lee, G. Disease-related modifications in tau affect the interaction between Fyn and Tau. J Biol Chem 280, 35119–35125 (2005).

86. Akache, B. et al. Anti-IgE Qb-VLP Conjugate Vaccine Self-Adjuvants through Activation of TLR7. Vaccines 4, 3 (2016).

87. Morris, R. Developments of a water-maze procedure for studying spatial learning in the rat. Journal of Neuroscience Methods 11, 47–60 (1984).

88. Vorhees, C. V. & Williams, M. T. Morris water maze: procedures for assessing spatial and related forms of learning and memory. Nat Protoc 1, 848–858 (2006).

89. Gallyas, F. Silver staining of Alzheimer’s neurofibrillary changes by means of physical development. Acta Morphol Acad Sci Hung 19, 1–8 (1971).

